# Focal adhesion kinase regulates early steps of myofibrillogenesis in cardiomyocytes

**DOI:** 10.1101/261248

**Authors:** Nilay Taneja, Abigail C. Neininger, Matthew R. Bersi, W. David Merryman, Dylan T. Burnette

**Affiliations:** Department of Cell and Developmental Biology, Vanderbilt University, Nashville, TN. USA. 37232; Department of Biomedical Engineering, Vanderbilt University, Nashville, TN. USA. 37232

## Abstract

Forces generated by myofibrils within cardiomyocytes must be balanced by adhesion to the substrate and to other cardiomyocytes for proper heart function. Loss of this force balance results in cardiomyopathies that ultimately cause heart failure. How this force balance is first established during the assembly of myofibrils is poorly understood. Using human induced pluripotent stem cell derived cardiomyocytes, we show coupling of focal adhesions to myofibrils during early steps of *de novo* myofibrillogenesis is essential for myofibril maturation. We also establish a key role for Focal adhesion kinase (FAK), a known regulator of adhesion dynamics in non-muscle cells, in regulating focal adhesion dynamics in cardiomyocytes. Specifically, FAK inhibition increased the stability of vinculin in focal adhesions, allowing greater substrate coupling of assembling myofibrils. Furthermore, this coupling is critical for regulating myofibril tension and viscosity. Taken together, our findings uncover a fundamental mechanism regulating the maturation of myofibrils in human cardiomyocytes.

## Introduction

The heart generates contractile force through the shortening of sarcomeres, consisting of myosin “thick” filaments forming sliding interactions with actin “thin” filaments. Actin filaments of adjacent sarcomeres are cross-linked by α-actinin-2 at Z-discs, which are important sites for intracellular signaling and mechanical stability of cardiomyocytes (Knoll et al., 2011; Kovacic-Milivojevic et al., 2001). The basic functional unit of contraction within a cardiomyocyte is the myofibril, comprised of a series of sarcomeres. In healthy cardiomyocytes, contractile forces generated by myofibrils are balanced by adhesive forces, specifically at cell-extracellular matrix (ECM) contacts (referred to as costameres), which transfer these forces to the ECM, and at cell-cell contacts (referred to as intercalated discs) (Liu et al., 2016; Samarel, 2005). Loss of this force balance can lead to detrimental phenotypes and disease states, such as dilated cardiomyopathy (DCM) (Dabiri et al., 2012; Samarel, 2005). Indeed, approximately 20% of causative DCM mutations are found in genes that encode either costameric or contractile proteins (Harvey and Leinwand, 2011; Hershberger et al., 2013). Unfortunately, how the adhesive and contractile systems assemble in cardiomyocytes is not well understood.

Previous work has shown that inhibiting contraction of cultured primary cardiomyocytes results in a loss of adhesions (Sharp et al., 1997; Simpson et al., 1993). Conversely, modulating the size of adhesions by varying substrate stiffness can modulate the contractile properties of the cardiomyocyte (Hersch et al., 2013; Jacot et al., 2008). This indicates that there is feedback between cardiomyocyte adhesion and contractile function. These studies investigated this relationship in cardiomyocytes with a pre-formed contractile apparatus. As such, the molecular pathways underlying this relationship during myofibril assembly remain poorly understood.

In culture, cardiomyocytes form focal adhesions (FAs) analogous to costameres which, *in vivo*, laterally anchor the membrane-proximal regions of Z-discs to the ECM (Pardo et al., 1983; Sharp et al., 1997). <u>F</u>ocal <u>a</u>dhesion <u>k</u>inase (FAK) is a key regulator of FA assembly and signal transduction in non-muscle cells (Parsons et al., 2000). In muscle cells, expression of dominant-negative FAK leads to disorganized myofibrils (Kovacic-Milivojevic et al., 2001). In primary cultured myocytes, increase in ventricular pressure and stretch can activate FAK through autophosphorylation (Domingos et al., 2002; Peng et al., 2006). This activation is necessary for sarcomere reorganization during the hypertrophic response (Chu et al., 2011; Pham et al., 2000). Furthermore, mice with cardiac-specific FAK knockout develop a DCM-like phenotype within 6 months (DiMichele et al., 2006; Peng et al., 2006). Therefore, FAK is an ideal candidate to regulate costamere assembly and myofibrillogenesis (Mansour et al., 2004).

While there is extensive evidence for the role of FAK in relaying mechanical stimuli to the contractile machinery, the cellular and molecular basis for this role are not known. Furthermore, the role of FAK during *de novo* myofibrillogenesis is not known. Investigating the role of FAK during this process has been limited by the lack of an appropriate model system to study *de novo* assembly of myofibrils and FAs. Classic work has utilized isolated primary rat, chick, and feline cardiomyocytes (Dabiri et al., 1997; Sharp et al., 1997; Simpson et al., 1993) to study myofibril and costamere assembly. However, these cells have technical limitations, such as low survival and poor amenability for genetic manipulation (Peter et al., 2016). In addition, these systems cannot be used to study *de novo* myofibril assembly, as they have pre-existing myofibrils.

In this study, we utilized induced pluripotent stem cell (iPSC)-derived human cardiomyocytes (hCMs) as a model system for *de novo* myofibril and adhesion assembly (Fenix et al., 2017). iPSC hCMs have robust survival and are amenable to genetic manipulation and exogenous protein expression, allowing for a unique opportunity to study myofibrillogenesis with unprecedented temporal and spatial resolution (Peter et al., 2016). Using this system, we show that FAK regulates the early steps of myofibrillogenesis. Genetic or pharmacological modulation of FAK regulates the assembly and dynamics of FAs in hCMs. We further show changes in adhesion size and stability affect the dynamics of thick and thin filament assembly, as well as the mechanical properties of myofibrils. Taken together, these experiments show a novel role for FAK, and for cell-ECM adhesion in general, in mediating the force balance between adhesion and contraction during myofibrillogenesis.

## Results and discussion

### Adhesion assembly and myofibrillogenesis are positively correlated

We used iPSC-derived human cardiomyocytes (hCMs) as a model system to study the simultaneous assembly of myofibrils and FAs. iPSC-derived hCMs form both myofibrils and FAs *de novo* over a period of 24 hours after plating. To visualize the assembly of myofibrils, we localized endogenous α-actinin-2, a protein that marks both muscle stress fibers (MSFs) and nascent myofibrils, characterized by small punctate structures known as Z-bodies, as well as mature myofibrils, characterized by longer Z-bodies known as Z-lines (Dabiri et al., 1997; Yang and Xu, 2012). At 6 hours post-plating, we noted that α-actinin-2 primarily localized as Z-bodies (Figure 1A), indicative of MSFs and nascent myofibrils. At 24 hours, we observed the formation of myofibrils characterized by longer Z-lines, with small Z-bodies restricted to the leading edge (Figure 1A).

Previous studies in spreading embryonic chick cardiomyocytes have shown that Z-bodies continuously incorporate into existing myofibrils (Dabiri et al., 1997). To investigate how the first Z-lines arise from Z-bodies in hCMs, we performed live imaging of α-actinin-2 during hCM spreading. We observed Z-bodies underwent retrograde movement and stitched to form the first Z-lines of myofibrils (Figure 1B). This retrograde movement continued for the duration of imaging (48 hours), concomitant with constant addition of nascent myofibrils into mature myofibrils (Supplementary Figure 1).

**Figure 1.**
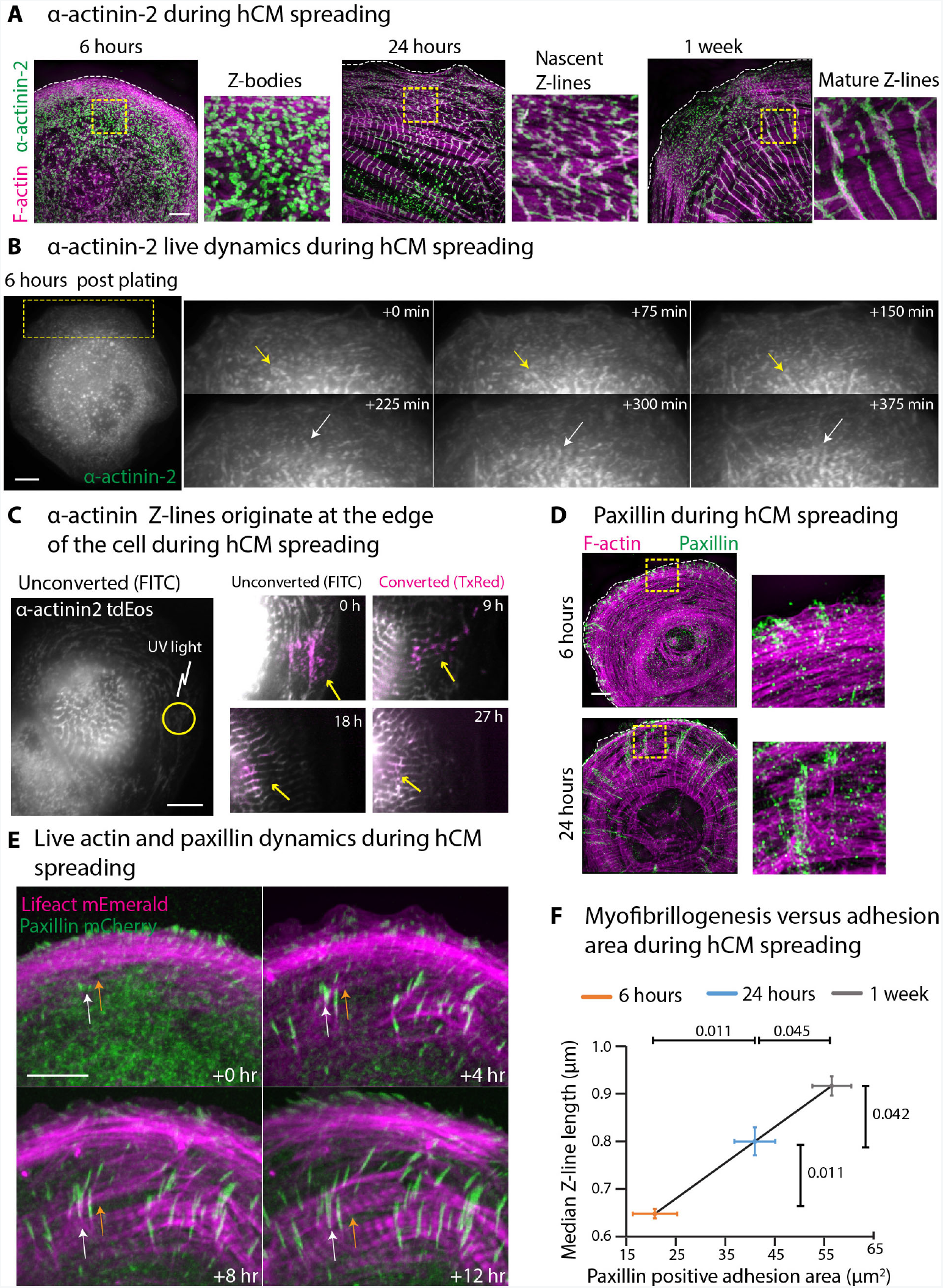
Adhesion maturation and myofibrillogenesis are positively correlated during hCM spreading. A) Representative SIM micrographs of hCMs spread for indicated times, visualizing F-actin and endogenous α-actinin-2. Insets: enlarged views of dotted yellow boxes. Dotted white lines: leading edge of the hCM. B) Live dynamics of α-actinin-2 during Hcm spreading. Inset: Time montage of dotted yellow box. 0 min corresponds to 6 hours post plating. Yellow and white arrow denote two separate Z-body stitching events. C) hCM expressing α-actinin-2 tdEos spread for 18 hours (corresponding to 0h in inset). Yellow circle represents ROI used for photo-conversion. Time montage shows unconverted α-actinin-2 (gray) overlaid with converted α-actinin-2 (magenta). Yellow arrows follow converted Z-bodies incorporating into Z-lines. D) Representative SIM micrographs of hCMs spread for 6 or 24 hours visualizing F-actin and endogenous paxillin. Insets show enlarged views of dotted yellow boxes. Dotted white lines show leading edge of the spreading hCM. E) Live dynamics of actin (magenta) and paxillin (green) during hCM spreading. 0 hours corresponds to 6 hours post plating. White arrows denote a maturing FA, and orange arrows show an assembling myofibril. F) Correlation between sum adhesion area and median Z-line length performed for time points indicated. Z-bodies (Y-axis): 6 hours-27877 Z-bodies from 80 cells over 3 experiments; 24 hours-33188 Z-bodies from 88 cells over 4 experiments; 1 week-62663 Z-bodies from 145 cells over 3 experiments. Adhesions (X-axis): n= 35, 38 and 28 cells over 4 independent experiments for 6 hours, 24 hours and 1 week, respectively. Error bars represent standard error of the mean. Scale bars: (A, D)-5 μm; (B, C, E)-10 μm.

We observed at later points in spreading (past 24 hours), myofibrils appeared to arise from large adhesion plaques parallel to the cell edge, that is, with sarcomeres aristing perpendicularly out of adhesion plaques, similar to what was recently claimed as “centripetal nucleation” (Supplementary Figure 1B) (Chopra et al., 2018). However, high temporal sampling of this process revealed that these myofibrils mainly arise by stitching of Z-bodies (Supplementary Figure 1C). To further confirm that Z-lines originate from Z-bodies at the edge of the cell, we utilized a photo-convertible probe, tdEos, that converts from green to red upon stimulation with UV light, to convert a population of α-actinin-2 Z-bodies at the edge of the hCM. We observed the converted population of α-actinin-2 in the Z-bodies persisted for up to 27 hours (Figure 1C). Thus, the α-actinin-2 that is incorporated into the Z-bodies at the edge persists long enough to be incorporated into mature Z-lines (Figure 1C, Supplementary Movie 1).

Interestingly, we observed that paxillin, a component of FAs, also localized in small punctate FAs at the cell edge at 6 hours post-plating (Figure 1D). Similar to α-actinin-2, at 24 hours, we observed the presence of larger FAs towards the cell body, and small punctate adhesions still localizing at the cell edge. Live imaging of paxillin together with actin during hCM spreading showed a small subset of FAs at the edge underwent retrograde movement and increased in size, or “matured” (Figure 1E, Supplementary Movie 2). This FA maturation was concomitant with the transition of nascent myofibrils to mature myofibrils. This strong temporal correlation suggested a feedback between myofibril and FA assembly.

To reduce bias and increase sample size, we developed an unbiased, computer-assisted automatic analysis that measures Z-line length and adhesion size in hundreds of cells during hCM spreading (Supplementary Figure 2). We used the length of Z-lines to represent the relative maturity of a developing myofibril, and total paxillin positive surface area to measure extent of adhesion. This measurement utilized images of spreading hCMs fixed and stained for endogenous α-actinin-2 and paxillin at 6 hours, 24 hours and 1 week. As suggested by our live data, we observed strong positive correlation between extent of adhesion and myofibril maturity at 6 hours, 24 hours, and up to 1 week post-plating (Figure 1F).

### Modulation of FAK activity affects myofibrillogenesis

Since adhesion assembly was positively correlated with myofibrillogenesis (Figure 1F), we asked if we could alter myofibrillogenesis by modulating FA stability. Focal adhesion kinase (FAK) is an integral regulator of focal adhesion turnover (Parsons et al., 2000; Webb et al., 2004). Since iPSC-derived cardiomyocytes are amenable to genetic manipulation, we utilized this advantage to knock down FAK using siRNA (si-FAK) and observed myofibrillogenesis in spreading hCMs. We obtained nearly 90% depletion of FAK protein levels using siRNA (Figure 2A). si-FAK hCMs showed a 40% reduction in adhesion area compared to scrambled controls (si-Scr) at 24 hours post-plating (Figure 2B). While these cells displayed paxillin-positive FAs, these adhesions completely lacked auto-phosphorylated FAK (Y397) (Figure 2A). We next predicted the reduction in adhesion area after si-FAK should impair myofibrillogenesis. Indeed, using our computer-assisted image analysis pipeline, we observed si-FAK hCMs had a shorter median Z-body length compared to si-Scr, indicative of being predominantly composed of MSFs and nascent myofibrils (Figure 2B).

**Figure 2.**
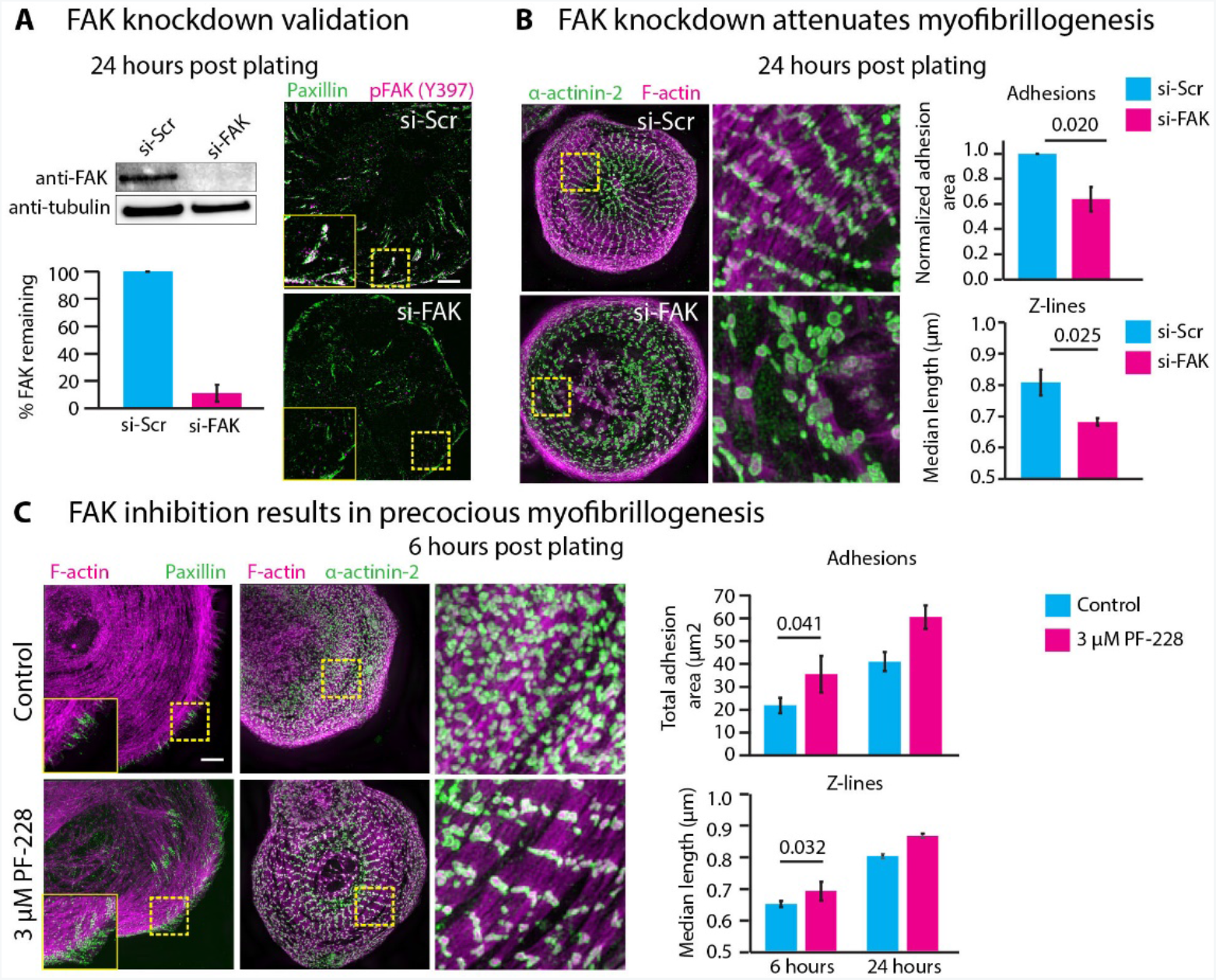
FAK regulates the rate of myofibril assembly. A) Validation of FAK knockdown using western blotting and immunofluorescence. For western blotting, tubulin was used as loading control for normalization. N=3 independent knockdowns. Representative SIM micrographs of scrambled control (si-Scr) and FAK depleted (si-FAK) hCMs spread for 24 hours. Endogenous paxillin and phospho-FAK (Y397) staining shows loss of phosphorylated FAK from adhesions and decrease in sum adhesion area. Inset: enlarged view of dotted yellow box. B) F-actin and endogenous α-actinin-2 staining at 24 hours shows overall reduction in Z-line length. Insets: enlarged view of dotted yellow box. Adhesions: sum paxillin area was quantified, as before. n=34 si-Scr and 29 si-FAK cells over 3 experiments. Z-lines: For si-Scr-19196 Z-bodies measured from 107 cells over 4 experiments. For si-FAK-28665 Z-bodies measured from 106 cells over 4 experiments. C) - Representative SIM micrographs of control versus PF-228 treated hCMs spread for 6 hours. Endogenous paxillin and F-actin staining shows increased adhesion area. Endogenous α-actinin-2 and F-actin staining shows longer Z-lines. Adhesion quantification: n=46 PF-228 treated cells over 4 experiments compared to control data set from Figure 1D. Z-line quantification: PF-228-21180 Z-bodies measured from 88 cells over 3 independent experiments, compared to control Z-line data set from Figure 1D. Error bars represent standard error of the means. Scale bars: 5 μm.

We next asked if increasing FA stability could accelerate the rate of myofibrillogenesis. Inhibition of the autophosphorylation of FAK with a small molecule inhibitor (PF-228) slows adhesion turnover without affecting formation during interphase and mitosis in non-muscle cells (Slack-Davis et al., 2007; Taneja et al., 2016; Webb et al., 2004). As expected, hCMs spreading in the presence of PF-228 showed increased adhesion area at 6 hours post-plating (Figure 2C, left). Surprisingly, in contrast to untreated cells at 6 hours post-plating, which do not contain any mature myofibrils, inhibition of FAK led to precocious myofibril assembly and increased median Z-body length (Figure 2C, right). This trend continued up to 24 hours post-plating (Figure 2C, right). Taken together, these experiments show that modulating FAK activity and therefore modulating the extent of adhesion alters the rate of myofibrillogenesis.

### FAK inhibition increases adhesion stability and slows retrograde movement of Z-bodies

We next wanted to investigate the mechanism underlying the precocious formation of myofibrils upon FAK inhibition. Since FAK inhibition led to increased adhesion area, we asked if FAK inhibition affected the dynamics of specific FA components. We again utilized the photo-convertible probe, tdEos, to measure turnover of adhesion components in control and FAK-inhibited cells. To compare our results with previous measurements in non-muscle cells using photo-conversion (Qin et al., 2015), we also measured adhesion dynamics in U2-OS cells. We first measured turnover of FAK itself, and observed a significant, ~2-fold increase in the half-life of FAK within FAs upon FAK inhibition (Figure 3A, Supplementary Figure 3). Notably, the half-life of FAK was significantly longer in control hCMs compared to U2-OS cells (Figure 3A).

**Figure 3.**
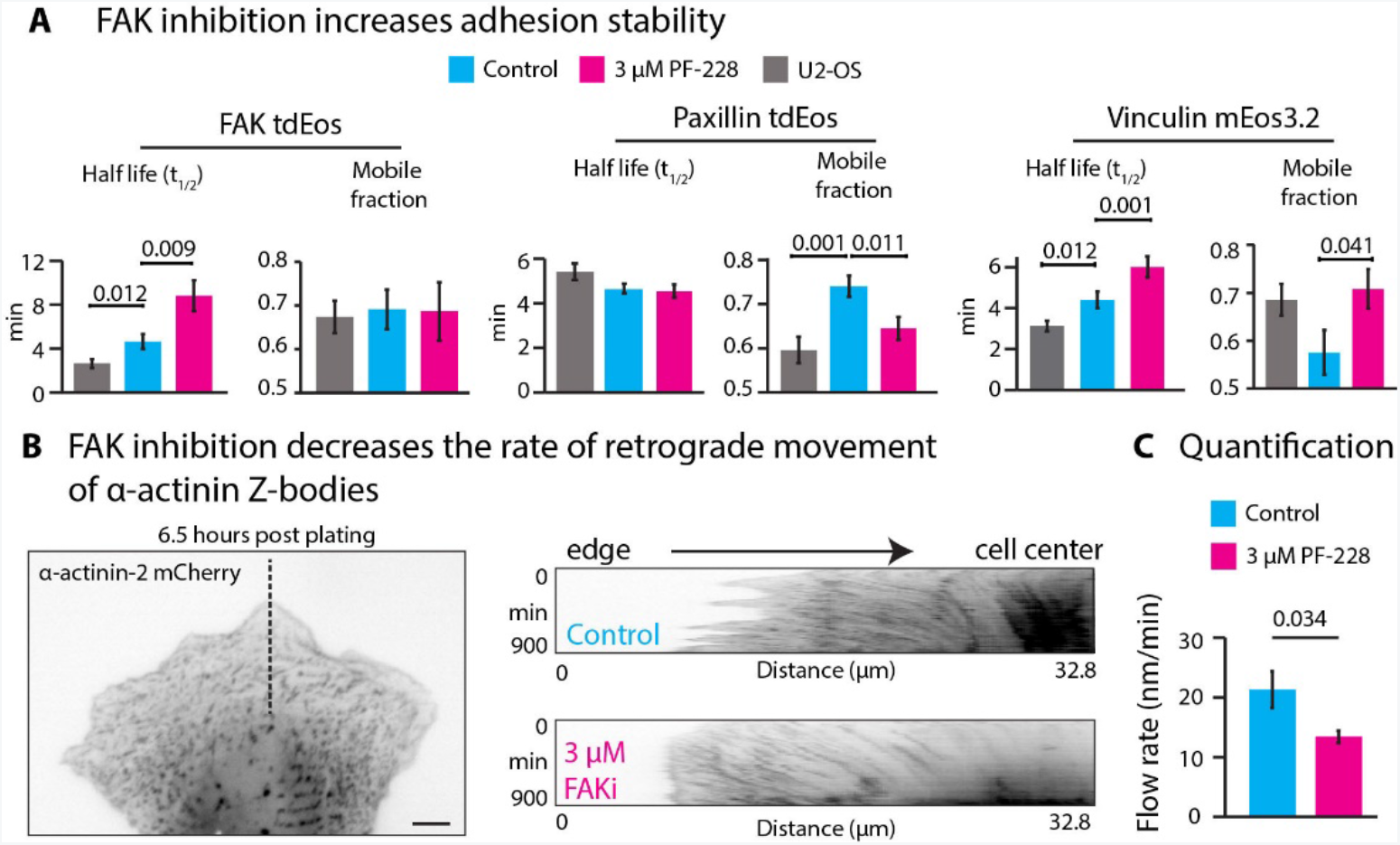
FAK inhibition results in adhesion stability and reduced retrograde flow rate. A) Half-lives and mobile fractions of FAK, vinculin and paxillin measured using photo-conversion in untreated U2-OS cells, untreated hCMs and PF-228 treated hCMs. FAK: n=15 U2-OS, 14 control hCM and 13 PF-228 treated hCM over 3 independent experiments. Vinculin: n=15 U2-OS, 14 control hCM and 15 PF-228 treated hCM over 3 independent experiments. B) Representative hCM expressing α-actinin-2 mCherry (inverted gray). Kymographs were generated using dotted black line to measure the rate of retrograde flow of Z-bodies. Representative kymographs for control versus PF-228 treated hCMs are depicted. E) Quantification of Z-body flow rates in control versus PF-228 hCMs. Control: 6 cells with 60 trajectories over 3 independent experiments. PF-228: 6 cells with 50 trajectories over 3 independent experiments. Error bars represent standard error of the means. Scale bars: 10 μm.

Since FAK is an integral scaffold protein that recruits and activates other proteins within FAs (Parsons et al., 2000), we hypothesized this stabilization of FAK should alter the dynamics of its interacting proteins. We found no change in the turnover rate of paxillin in control hCMs upon FAK inhibition or in comparison to U2-OS cells; however, we observed a small but significant decrease in the dynamic pool, or mobile fraction, upon FAK inhibition, in agreement with increased adhesion area (Figure 3A). Vinculin is an integral component of FAs that is known to act as the “clutch” that transduces force from the ECM to the actin cytoskeleton (Kanchanawong et al., 2010; Thievessen et al., 2013). Interestingly, vinculin showed a similar trend as FAK; the half-life of vinculin was significantly longer in control hCMs versus U2-OS cells, and inhibition of FAK in hCMs further increased the half-life (Figure 3A). Therefore, FAs in hCMs are more stable compared to non-muscle cells, and FAK inhibition further increases hCM adhesion stability.

Previous studies show that during non-muscle cell migration, adhesions can act to slow retrograde movement by increased substrate coupling of stress fibers to facilitate leading edge advance (Burnette et al., 2011; Thievessen et al., 2013). This led us to hypothesize that larger adhesions upon FAK inhibition would similarly slow MSFs as they undergo retrograde movement. To test this, we measured retrograde movement of Z-bodies in α-actinin-2-mCherry transfected hCMs and found that FAK inhibition indeed decreased the rate of retrograde movement of α-actinin-2 Z-bodies (Figure 3B-C). These experiments strongly suggest FAK inhibition stabilizes adhesion components, which allows greater coupling of myofibrils with the ECM, thus slowing down the retrograde movement of MSFs as they flow backwards from the edge to assemble into nascent myofibrils.

### FAK inhibition affects thin filament and thick filament components

We next asked how the slowing down of retrograde movement upon FAK inhibition affected myofibrillogenesis. We hypothesized that slower movement, along with greater substrate coupling, should allow other sarcomeric components to incorporate into the myofibril. The giant protein titin loads early on in myofibrillogenesis and is the first marker of nascent myofibrils (Rhee et al., 1994). To visualize titin, we localized the N-terminal domain of titin, which localizes at Z-lines (Rhee et al., 1994). In control hCMs at 6 hours post-plating, titin appeared as punctate bodies distributed throughout the cell, with some limited organization at the cell center (Figure 4A, left). Upon FAK inhibition, we observed a dramatic increase in organization of titin, appearing to flank the bright actin foci that correspond to Z-bodies (Figure 4A). To quantify these changes in titin organization, we measured the distance from the cell edge where titin first appears and found that titin loaded closer to the cell edge in FAK-inhibited cells (Figure 4B).

**Figure 4.**
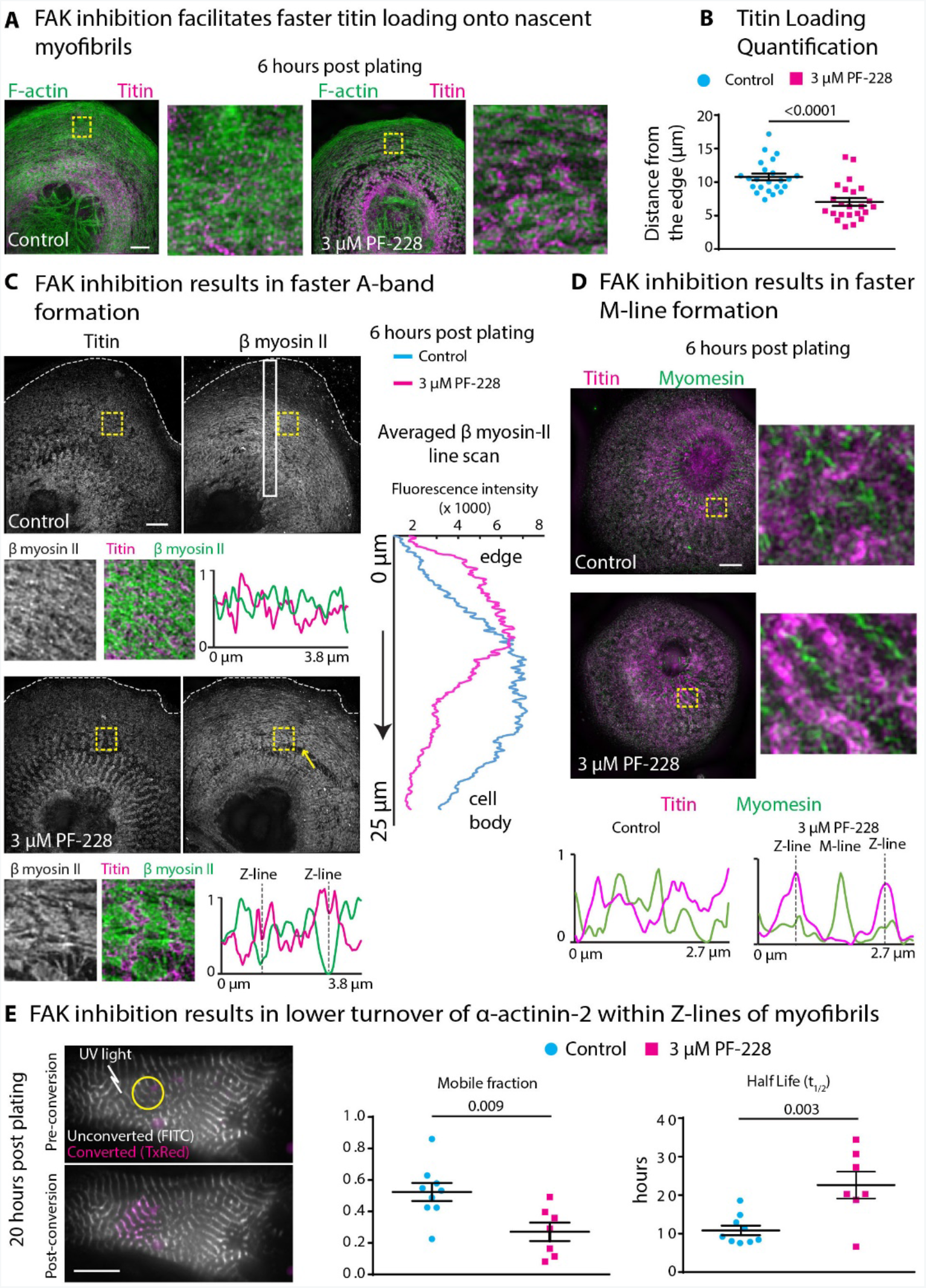
FAK inhibition results in faster A-band and M-line formation and increased Z-line stability. A) Representative SIM micrographs of hCMs allowed to spread for 6 hours showing endogenous titin and F-actin in control versus PF-228 treatment. Inset: Enlarged view of dotted yellow box. B) Measurement of distance from the edge for the appearance of first titin “rings” around Z-bodies. n=23 cells each for control and PF-228 over 3 experiments. C) Representative SIM micrographs of hCMs allowed to spread for 6 hours showing endogenous β-myosin II and titin in control versus PF-228 treatment. Inset: Enlarged view of dotted yellow box and representative line scans. Note sarcomeric organization in PF-228 treated hCMs. White boxes were used to generate an averaged line scan for β-myosin II in control (n=11 cells, 3 experiments) and PF-228 (n=10 cells, 3 experiments) show increased β-myosin II incorporation into myofibrils upon PF-228 treatment. D) Representative SIM micrographs of hCMs allowed to spread for 6 hours showing endogenous myomesin and titin in control versus PF-228 treatment. Insets: Enlarged view of dotted yellow box and representative line scans. Note sarcomeric organization in PF-228 treated hCMs. E) Representative hCM at 16 hours expressing α-actinin-2 tdEos. Yellow circle represents photo-converted (show in magenta) area. Bar graphs compare half-life and mobile fraction upon PF-228 treatment. Error bars represent standard error of the mean. Scale bars: (A, C, D)-5 μm; (E)-10 μm.

The role of titin in the sarcomere is to stabilize the thick and thin filaments and act as a spacer to center the thick filament between the thin filaments (Anderson and Granzier, 2012; Rhee et al., 1994; Sanger et al., 2005). Therefore, we hypothesized that β-myosin should organize around titin observed upon FAK inhibition. In control hCMs 6 hours post-plating, β-myosin appeared as continuous filaments, rather than as the stack of filaments characteristic of the “A-band”, as reported previously (Rhee et al., 1994). Consistent with our prediction, upon FAK inhibition, we observed precocious formation of A-bands around titin, with an overall increase in the amount of β-myosin closer to the edge (Figure 4C, line scans of average intensity of β-myosin). Line scans of titin and β-myosin (Figure 4C, insets) showed a clear sarcomeric organization in FAK-inhibited cells, but not in control cells. Furthermore, formation of an M-line is the latest step in forming a mature myofibril (Ehler et al., 1999). In FAK-inhibited cells, long continuous myomesin bands alternated with titin, while this organization was not observed in control hCMs (Figure 4D). Taken together, these results show that FAK inhibition leads to precocious loading of both thin filament and thick filament components, accelerating the maturation of myofibrils.

Given that FAK inhibition leads to increased incorporation of sarcomeric proteins, we asked whether this leads to increased stability of myofibrils. To test this, we measured the turnover of α-actinin-2 within Z-lines of mature myofibrils using photo-conversion. In control hCMs, we noted limited turnover of α-actinin-2 in Z-lines, with a half-life of ~10 hours. Upon FAK inhibition, we observed a significant increase in the half-life as well as a decrease in the mobile fraction (Figure 4E). This increased Z-line stability, along with precocious incorporation of titin and β-myosin then led us to test whether this has an influence on mechanical properties of these myofibrils.

### FAK inhibition increases myofibril viscosity

We therefore characterized the mechanical properties of individual myofibrils based on the time-dependent end separation of myofibrils following focused laser weakening (Kumar et al., 2006). Using a modified edge detection algorithm, separation profiles were measured in myofibril kymographs from either control or FAK-inhibited hCMs (red traces; Figure 5A). Using a standard Kelvin-Voigt viscoelastic model, characterized by an asymptotic distance *D*_a_ that is correlated with elastic energy dissipation and a relaxation time constant τ that reflects the ratio of viscosity to elasticity (cf. Figure 5B) (Kassianidou et al., 2017), we were able to obtain good fits to both control and FAK-inhibited separation profiles (*R*^2^>0.99; Figure 5C). In addition to viscoelastic parameters, we used best-fit curves to calculate the instantaneous separation velocity, an estimate of myofibril tension (Peralta et al., 2007), and the linear separation velocity, a measure of overall retraction speed.

**Figure 5.**
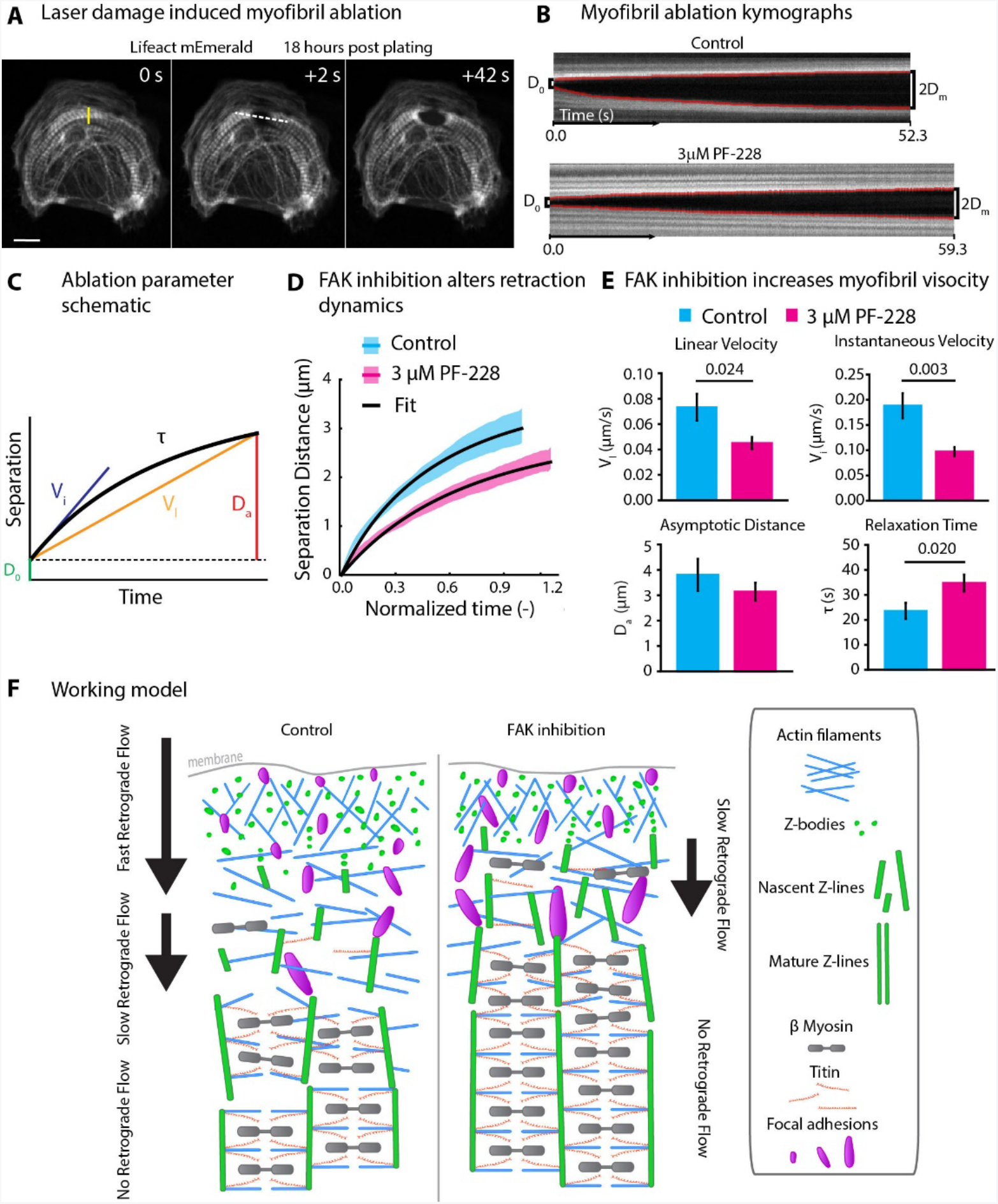
FAK inhibition alters myofibril viscous properties. A) Representative time montage of hCM expressing Lifeact mEmerald. The yellow line was used as the ROI for ablation using laser damage. B) Representative kymographs from control versus PF-228 hCMs generated using lines drawn parallel to the retracting myofibril, such as dotted white line in (A). Measurements from kymographs were made using a custom MATLAB script (see Methods) for detecting the edges of the retracting myofibrils (solid red lines). D_0_: Initial separation; D_m_: Maximal separation of each side of the retracting myofibril. C) Schematic of parameters obtained from the fitted retraction curve (solid black curve). V_i_-Instantaneous velocity; τ-Relaxation time; V_l_-linear velocity; D_a_-Asymptotic distance. D) Averaged retraction curves of hCMs comparing control versus PF-228 treated hCMs. Shaded blue and magenta regions show standard error of the means. n=18 control and 24 PF-228 treated hCMs over 3 independent experiments. E) Comparison of linear velocity, instantaneous velocity, asymptotic distance and relaxation time (as described in (C)) in control versus PF-228 hCMs. Error bars represent standard error of the means. Scale bar: 10 μm. F) Working model for role of FAK during myofibrillogenesis.

Despite similar levels of asymptotic separation (i.e., energy dissipation), FAK-inhibited myofibrils showed a more viscous separation response than control myofibrils (Figure 5D). Indeed, increased viscous relaxation times contributed to a reduction in linear separation velocity due to the increased time required for maximum separation. Given that myofibril viscosity is largely dictated by actin-titin interactions (Kulke et al., 2001; Linke and Fernandez, 2002), it is not surprising that FAK-inhibited cells show increases in both titin incorporation and myofibril viscosity. Together, these results indicate that FAK regulates the mechanical properties of mature myofibrils in a manner that is consistent with our prior observations of reduced retrograde flow and faster myofibril component incorporation.

### Model of myofibrillogenesis regulation by adhesion stability

In this study, we describe the role of FAK in regulating the early steps of myofibrillogenesis. We show that adhesion assembly was strongly positively correlated with myofibrillogenesis at early time points (6 and 24 hours post-plating) and continued up to 1 week (Figure 1). Knockdown or inhibition of FAK led to decreased and increased cell adhesion, respectively, resulting in delayed and accelerated myofibrillogenesis, respectively (Figure 2). We further show FAK inhibition led to increased adhesion stability, leading to increased coupling of myofibrils to the substrate, concomitant with a decrease in retrograde flow rate of α-actinin-2 Z-bodies (Figure 3). This led to faster incorporation of sarcomeric components titin and β-myosin, accompanied by increased stability of Z-lines (Figure 4). Ultimately, this leads to increased myofibril viscosity (Figure 5).

The role of FAK has been previously investigated in non-muscle cells in the context of cell migration and mitosis (Parsons et al., 2000; Taneja et al., 2016). In muscle cells, FAK localizes to costameres and intercalated discs, and its role in FAK/Src signaling is well appreciated (Franchini et al., 2009; Yi et al., 2003). However, the mechanism(s) by which FAK regulates myofibrillogenesis was previously unknown. In this study, we show that FAK regulates adhesion stability in human cardiomyocytes, which led us to propose a novel role for cell-ECM adhesion in the early steps of myofibrillogenesis (Figure 5F). As Z-bodies move toward the cell body, a subset of FAs at the edge engage muscle-specific stress fibers, resulting in slower retrograde flow. These FAs increase in size while also undergoing retrograde flow; this suggests they experience increased tension, since FAs mature upon experiencing actomyosin generated tension (Oakes and Gardel, 2014). Our finding that titin and β-myosin incorporate faster upon FAK inhibition suggests a bidirectional mechanical coupling between FA maturation and increased force production through the maturation of myofibrils.

The model we are proposing in Figure 5F is an extension of a model we recently proposed based on data that clearly shows that myofibrils arise from muscle-specific stress fiber precursors (Fenix et al., 2017). These myofibril precursors are assembled parallel to the edge of the cell and move away from the cell through retrograde flow while sarcomeres elaborate along their lengths. In particular, Z-lines of the new assembling sarcomeres are perpendicular to the edge (Fenix et al., 2017) and parallel to the long axis of mature FAs (Figure 5F). However, a recent study has proposed an alternative model of myofibrillogenesis, where myofibrils nucleate directly from FAs and have their Z-lines positioned with the opposite orientation as our data shows (Chopra et al., 2018). That is, sarcomeres are imagined to be streaming out of the adhesions with their Z-lines positioned parallel to the edge of the cell and perpendicular to the long axis of adhesions. Undoubtedly, this model and the one we proposed in Figure 5F are fundamentally different, and we sought to determine the origin of these differences.

Almost all of the hCMs presented by Chopra et al. appear to have similar morphologies to our cells. Indeed, the Z-lines in these cells are positioned perpendicular to the edge as we propose. However, the hCMs used by Chopra et al had been plated for at least 24 hours and, as such, already had assembled myofibrils. This led us to examine our hCMs that were spread for at least 24 hours. Indeed, we found a few myofibrils that did have a morphology that could match that of the model proposed by Chopra et al. (Supplementary Figure 1). Indeed, imaging these myofibrils with low temporal sampling as presented by Chopra et al. could be interpreted as the Z-lines originating from the adhesion. However, higher temporal sampling clearly revealed that the Z-lines did not originate from the adhesion (Supplementary Figure 1C). As such, we propose that it is unlikely that the model proposed by Chopra et al. is an accurate representation of sarcomere assembly and if it indeed occurs, it is a rare event. Instead, our results show that myofibrils arise from immature myofibrils parallel to the leading edge of the cell, that is, with perpendicularly arising Z-lines from coalescing Z-bodies, and that adhesions serve to act as a ‘clutch’ that allows increased coupling of myofibrils to the substrate, similar to the coupling of “actin arc” stress fibers and adhesions in non-muscle cells (Burnette et al., 2011; Thievessen et al., 2013; Welch et al., 1997).

Cell-cell adhesion is another source of balancing forces generated by myofibrils (Liu et al., 2016). However, they are unlikely to play a significant role in myofibril assembly. Indeed, other groups have shown cell-ECM adhesion sites are responsible for majority of traction force generation during early cardiomyocyte spreading; as the cell continues to spread and makes cell-cell contact, load-bearing sites are transferred to cell-cell contact sites (McCain et al., 2012). As such, it is interesting to note that FA proteins such as FAK and vinculin also localize to intercalated discs (Koteliansky and Gneushev, 1983; Yi et al., 2003). Determining the mechanisms driving the localization of these proteins to sites of increased load should be the focus of future studies.

Finally, we establish iPSC-derived hCMs as a powerful model system for adhesion and myofibril assembly. iPSC-derived hCMs have distinct advantages over using primary cardiomyocytes, including robust survival, ease of genetic manipulation and being amenable to high resolution imaging. However, caution must be exercised in selecting the time window for the process being investigated, the spatial and temporal sampling required and using quantitative methods. We expect this system to be useful for studying multiple aspects of myofibril assembly and maintenance.

## Acknowledgements

We would like to thank the lab of Irina Kaverina, Vanderbilt University, for the gift of anti-FAK antibody. We would also like to thank the Bryan Millis and Sean Schaffer at the Nikon Center of Excellence and the Cell Imaging Shared Resource, Vanderbilt University for access to microscopes, software and technical support. This work was supported by a MIRA Grant R35GM125028-01 from NIGMS to D.T.B, a MIRA Grant R35HL135790 to W.D.M., and Vanderbilt University School of Medicine Program in Developmental Biology (Training grant T32-HD007502 to A.C.N). The authors declare no competing financial interest.

## Author Contributions

N.T. and D.T.B conceived the study. A.C.N. and N.T. performed the experiments. A.C.N, N.T. and M.R.B. analyzed the data. A.C.N., N.T., M.R.B and D.T.B wrote the manuscript. A.C.N developed the code to analyze photo-conversion, Z-line and focal adhesion datasets. M.R.B and W.D.M implemented the analysis of myofibril retraction profiles. All authors read and commented on the manuscript. The order of the co-first authors was determined by a coin flip.

## Materials and Methods

### Cell culture and chemicals

U2-OS (HTB-96; American Type Culture Collection, Manassas, VA) cells were cultured in 25-cm^2^ cell culture flasks (25-207; Genessee Scientific Corporation, San Diego, CA) in growth medium comprising DMEM (10-013-CV; Mediatech, Manassas, VA) containing 4.5 g/L L-glutamine, D-glucose, and sodium pyruvate and supplemented with 10% fetal bovine serum (F2442; Sigma-Aldrich, St. Louis, MO). iPSC-derived human cardiomyocytes (hiCMs, Cellular Dynamics, Madison, WI) were cultured as per manufacturer’s instructions in proprietary manufacturer provided cardiomyocyte maintenance medium (CMM-100-120-001, Cellular Dynamics, Madison, WI) in polystyrene 96-well cell culture plates.

Cells were cultured at 37°C and 5% CO_2_. For re-plating hCMs onto glass substrates, cells were washed 2x with 100 μL 1x PBS with no Ca^2n+^/Mg^2n+^ (PBS*, 70011-044, Gibco, Grand Island, NY). PBS* was completely removed from hiCMs and 40 μL 0.1% Trypsin-EDTA with no phenol red (15400-054, Gibco, Grand Island, NY) was added to hCMs and placed at 37° C for 2 minutes. Following incubation, the culture dish was washed 3x with trypsin inside well, rotated 180 degrees, and washed another 3x. Trypsinization was then quenched by adding 160 μL of culture media and total cell mixture was placed into a 1.5 mL Eppendorf tube. Cells were centrifuged at 1000xg for 5 minutes, and the supernatant was aspirated. Cells were then re-suspended in 200 μL of culture media and plated on 35 mm dishes with 10 mm glass bottom (D35-10-1.5-N; CellVis, Sunnydale, CA) pre-coated with 10 μg/mL fibronectin (#354008, Corning) for 1 hr at 37°C.

FAK inhibitor PF-228 (PZ0117) was purchased from Sigma. Alexa Fluor 488-phalloidin (A12379), Alexa Fluor 568-phalloidin (A12380), Alexa Fluor 488-goat anti-mouse (A11029), Alexa Fluor 488-goat anti-rabbit (A11034), Alexa Fluor 568-goat-anti-rabbit (A11011), and Alexa Fluor 568-goat anti-mouse (A11004) antibodies were purchased from Life Technologies (Grand Island, NY).

Mouse anti-Paxillin (1:200, 610051) and mouse anti-FAK (1:500, 610088) antibodies were purchased from BD Biosciences. Rabbit anti-FAK Y357 (1:200, ab81298) was purchased from Abcam. Mouse anti-α actinin-2 (1:200, A7811) was purchased from Sigma Aldrich. β-myosin II (1:1, A4.1025), Titin (9D10) and myomesin (1:1, mMAC B4) antibodies were purchased from the Developmental Studies Hybridoma Bank (University of Iowa). Primary antibody conjugation, if required, was performed using Mix-n-Stain kits (#92233) purchased from Biotium (Fremont, CA) according to the instructions provided by the manufacturer.

Paraformaldehyde (15710) was purchased from Electron Microscopy Sciences (Hatfield, PA). Triton X-100 (BP151100) was purchased from Fischer Scientific (Suwanee, GA).

### Fixation and immunostaining

Cells were fixed with 4% paraformaldehyde (PFA) in PBS at room temperature for 20 min and then extracted for 5 min with 1% Triton X-100 and 4% PFA in PBS as previously described (Burnette et al., 2014). For actin visualization, phalloidin 488 or 568 in 1× PBS (15 μl of stock phalloidin per 200 μl of PBS) was used for 2 hours at room temperature. For immunofluorescence experiments, cells were blocked in 10% bovine serum albumin (BSA) in PBS, followed by antibody incubations.

For visualizing β-myosin and titin, a live cell extraction was performed to remove cytoplasmic background as described previously (Burnette et al 2011). Briefly, a cytoskeleton-stabilizing live-cell extraction buffer was made fresh containing 2 ml of stock solution (500 mM 1,4-piperazinediethanesulfonic acid, 25 mM ethylene glycol tetraacetic acid, 25 mM MgCl2), 4 ml of 10% polyoxyethylene glycol (PEG; 35,000 molecular weight), 4 ml H2O, and 100 μl of Triton X-100, 10 μM paclitaxel, and 10 μM phalloidin. Cells were treated with this extraction buffer for 1 min, followed by a 1-min wash with wash buffer (extraction buffer without PEG or Triton X-100). Cells were then fixed with 4% PFA for 20 min, followed by antibody incubations.

VectaShield with DAPI (H-1200, Vector Laboratories Inc., Burlingame, CA) was used for mounting.

### Protein Expression

For protein expression in hCMs, 200 ng plasmid and 0.4 μl ViaFect (E498A, Promega, Madison WI) were diluted in a total of 10 μl of Opti-MEM (ThermoFisher, Waltham, MA) and added to a single well of hiCMs in a 96 well plate. The transfection was allowed to go overnight (~16 hrs). U2OS cells were transfected using Fugene 6 (E2691, Promega) as per instructions provided by the manufacturer.

### Plasmids

mCherry-Alpha-Actinin 2 (54974), tdEos-Alpha-Actinin 2 (57577), tdEos-Paxillin (57653), tdEos-PTK2 (57659), mEos3.2-Vinculin (66952), mEmerald Lifeact (54148) and mCherry-Paxillin (55114) were purchased from Addgene (Cambridge, MA).

### Knockdown of FAK using si-RNA

Knockdown of FAK (PTK2) was performed using Accell SmartPool siRNA (PTK2: E-003164-00-0005; scrambled control: D-001910-10-05) purchased from GE Dharmacon. Experiments were performed in 96-well culture plates, using the Transit-tKO (MIR2154, Mirus Bio) transfection reagent using instructions provided by the manufacturer. Three consecutive rounds of knockdown were performed to completely deplete FAK protein levels. Following knockdown, cells were re-plated onto glass substrates for immunofluorescence or lysed for western blotting.

### Western Blotting

Gel samples were prepared by mixing cell lysates with LDS sample buffer (Life Technologies, #NP0007) and Sample Reducing Buffer (Life Technologies, #NP00009) and boiled at 95°C for 5 minutes. Samples were resolved on Bolt 4-12% gradient Bis-Tris gels (Life Technologies, #NW04120BOX). Protein bands were blotted onto a nylon membrane (Millipore). Blots were blocked using 5% NFDM (Research Products International Corp, Mt. Prospect, IL, #33368) in TBST. Antibody incubations were also performed in 5% NFDM in TBST. Blots were developed using the Immobilon Chemiluminescence Kit (Millipore, #WBKLS0500).

### Structured Illumination Microscopy

SIM imaging and processing was performed on a GE Healthcare DeltaVision OMX equipped with a 60×1.42 NA Oil objective and sCMOS camera.

### Fluorescence, photo-conversion, and live-cell microscopy

High-resolution wide-field fluorescence images were acquired on a Nikon Eclipse Ti equipped with a Nikon 100× Plan Apo 1.45 numerical aperture (NA) oil objective and a Nikon DS-Qi2 CMOS camera. Photo-conversion was accomplished by blocking the Eclipse Ti’s field diaphragm using a sheet of foil with a small ROI created by puncturing with a microneedle, and using a 4’,6-diamidino-2-phenylindole filter cube to shine UV light on the structure of interest for 6 s as previously described (Fenix et al., 2016). This was immediately followed by opening the field diaphragm and imaging with FITC and TxRed filter cubes every 2, 3, 15, or 180 minutes depending on the protein expressed. For live imaging, cells were maintained at 37°C with 5% CO2 using a Tokai Hit stage incubator.

### Laser damage induced myofibril ablation

hCMs were transiently transfected with Lifeact mEmerald and re-plated on glass growth substrate as described above and allowed to spread for 18 hours. Cells were either untreated or treated with 3 μM PF-228 for 2 hours. Myofibril ablation was performed on an inverted Nikon Eclipse Ti-E inverted microscope equipped with a Yokogawa CSU-X1 spinning disk head, 1.4 NA 60X oil objective, Andor DU-897 EMCCD and a dedicated 100 mW 405 diode ablation laser, generously provided by the Nikon Centre of Excellence at Vanderbilt University. The ablation was laser was guided along a 4 μm long stimulation line using a miniscanner controlled in Nikon Elements software. A pixel dwell time of 500 μs, at 50% laser power for 1 second was used. Subsequent image stacks were acquired at an overall frame rate of 6 fps for a minimum of 60 seconds. To measure instantaneous recoil velocities, a 20% laser power was used. This allowed us to weaken the myofibrils and subsequently capture the initial recoil at 6 fps.

### Data Quantification

To measure Z-line length in hCMs (Figure 1C-D, Figure 2), Z-sections were acquired at 200 nm intervals. Images were deconvolved using Richardson-Lucy 3D deconvolution on Nikon Elements software. The α-actinin-2 bodies were thresholded manually in Nikon Elements using a 3x clean and excluding all bodies smaller than 0.2 μm in length. The binaries were combined in 3D using Elements software, and the lengths of the major axis of each body were exported for the calculation of median Z-line length. Size exclusion and calculation of Z-line lengths were performed using MATLAB.

To measure sum adhesion area of hCMs (Figure 1C-D, Figure 2), one single Z-slice of a reconstructed structured illumination image in the plane of focus of the adhesions was selected. Focal adhesions were thresholded and sum area was measured in Fiji, using a size minimum exclusion criterion of 0.1 μm^2^.

For photo-conversion experiments (Figure 3), time lapse montages of the converted channel (TxRed) were aligned using the StackReg plugin in Fiji. A rectangular ROI was drawn around the photo-converted area, and mean intensity over time was calculated using the Multi Measure function in the ROI manager of Fiji. Background correction was performed using a similar ROI placed in the background for each corresponding image. Additionally, we also subtracted any fluorescence signal present prior to photo-conversion. To do this, an image of the TxRed channel was recorded before photo-conversion, and mean intensity of the same ROI was measured, background corrected and subtracted from each post-photo-conversion image. To account for any bleaching the sample undergoes, we also performed photo-conversion experiments on fixed cells at the same temperature and media conditions and quantified with the same method. We created a MATLAB application to fit photo-conversion curves and calculate decay time and mobile fraction of each protein. The intensity measurements over time for each converted area were normalized to 1. An averaged bleaching curve was created, and the data was corrected for any bleaching. The script fit the curves of each converted area to a linear curve, a first-degree exponential, and a second-degree exponential using the curve-fitting toolbox and determined which curve was the best fit using a chi-squared test. The half-life of each curve was calculated as the time taken for half the maximal fluorescence loss. The mobile fraction was calculated as the amount of fluorescence loss as a fraction of the total fluorescence of the post-stimulated region. A curve showing an averaged fit across all regions converted is generated, showing mean +/- SEM.

For measurement of the distance of titin to the edge (Figure 4B), we measured the length of a line from the edge to the first titin “ring” in Fiji. The researcher analyzing the data was blinded to treatment groups. To produce the averaged β-myosin line scan (Figure 4C), we measured averaged intensity of a 100 pixel-wide line 20 μm long starting at the cell edge in Fiji, directed perpendicularly to the leading edge towards the cell center.

To quantify myofibril mechanical properties, individual myofibrils were subjected to laser weakening in order to induce separation. Following visible separation, kymographs were drawn along the separation axis in order to visualize the time-varying end-to-end distance. Using a custom edge detection algorithm based on the Canny filter, edge profiles were defined based on gradients in the kymograph image and the separation distance *L*(t) was computed as one-half of the distance between the upper and lower edge profiles at each frame (red curves, Figure 5A). Separation profiles were then fit to a standard Kelvin-Voigt viscoelastic solid model of the form

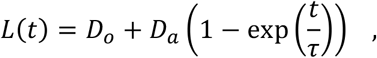

in order to identify two viscoelastic parameters: the asymptotic distance *D*_a_ and the relaxation time constant τ Note that the asymptotic distance parameter has been shown to be correlated with elastic energy dissipation, whereas the relaxation time constant is representative of the ratio of viscosity to elasticity in the material. Hence, increasing τ indicates a more viscous response and vice-versa. Additionally, any initial damage induced by the laser was accounted for by defining an offset distance *D_0_* Values of *D_0_* were not statistically different between control and FAK-inhibited cells.

### Statistics

Statistical significance of sum adhesion area and median Z-body length was determined using unpaired two-tailed Student’s T-tests performed in MATLAB or Excel. Each experiment was performed a minimum of 3 times and the mean of the independent experiments and standard error of the mean (SEM) are displayed. For photo-conversion, titin quantification and laser ablation experiments, each adhesion or cell is counted as a data point and each experiment was performed a minimum of 3 times.

## Supplementary Data

### Movie legends

### Supplementary Movie 1-

hCM expressing α-actinin-2-tdEos (magenta) to mark Z-bodies and Z-lines. A region of α-actinin-2 Z-bodies is converted (green) with UV light and observed undergoing retrograde flow and stitching into Z-lines of myofibrils.

Movie Length: 7 hours. Horizontal and vertical dimensions, 36 μm and 19 μm, respectively.

### Supplementary Movie 2-

hCM expressing Life-Act mEmerald (green) and paxillin-mCherry (magenta) spreading, showing a small subset of FAs (shown with paxillin) at the cell edge undergo retrograde movement toward the cell body and an increase in size (‘maturation’).

Movie Length: 10 hours. Horizontal and vertical dimensions, 64 μm and 65 μm, respectively.

---

**Supplementary Figure 1.**
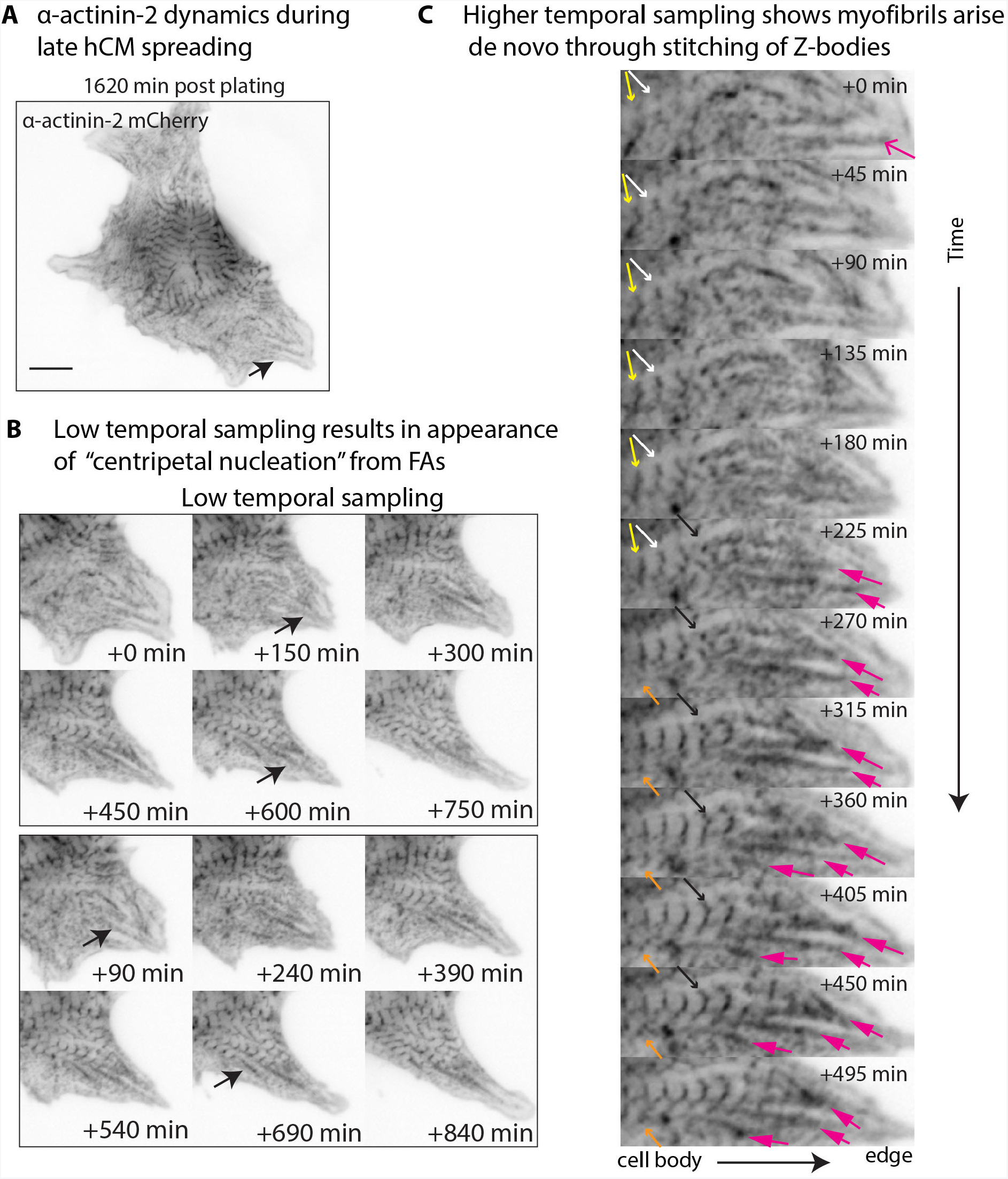
Myofibrils do not arise by centripetal nucleation from FAs but by lateral stitching of Z-bodies. A) hCM expressing a-actinin-2 mCherry at 27 hours post plating. B) Time montages of region correponding to black arrow displayed at 150 min intervals. Note the appearance of centripetal nucleation from long FA (black arrows). C) Higher temporal sampling shows several examples of the appearance of Z-bodies (orange, black, yellow and white arrows) far away from the initial FA (magenta arrows). Subsequent time frames show the myofibril arising by stitching of Z-bodies (white and black arrows). The adhesions (magenta arrows) subsequently grow brighter as the myofibril incorporates more Z-bodies. Note how Z-body in time frame 0 min (white arrow) is already elongated before the Z-body to the right (yellow arrow) appearing without any observeable from the adhesion at the edge. Scale bar: 10 µm.

**Supplementary Figure 2.**
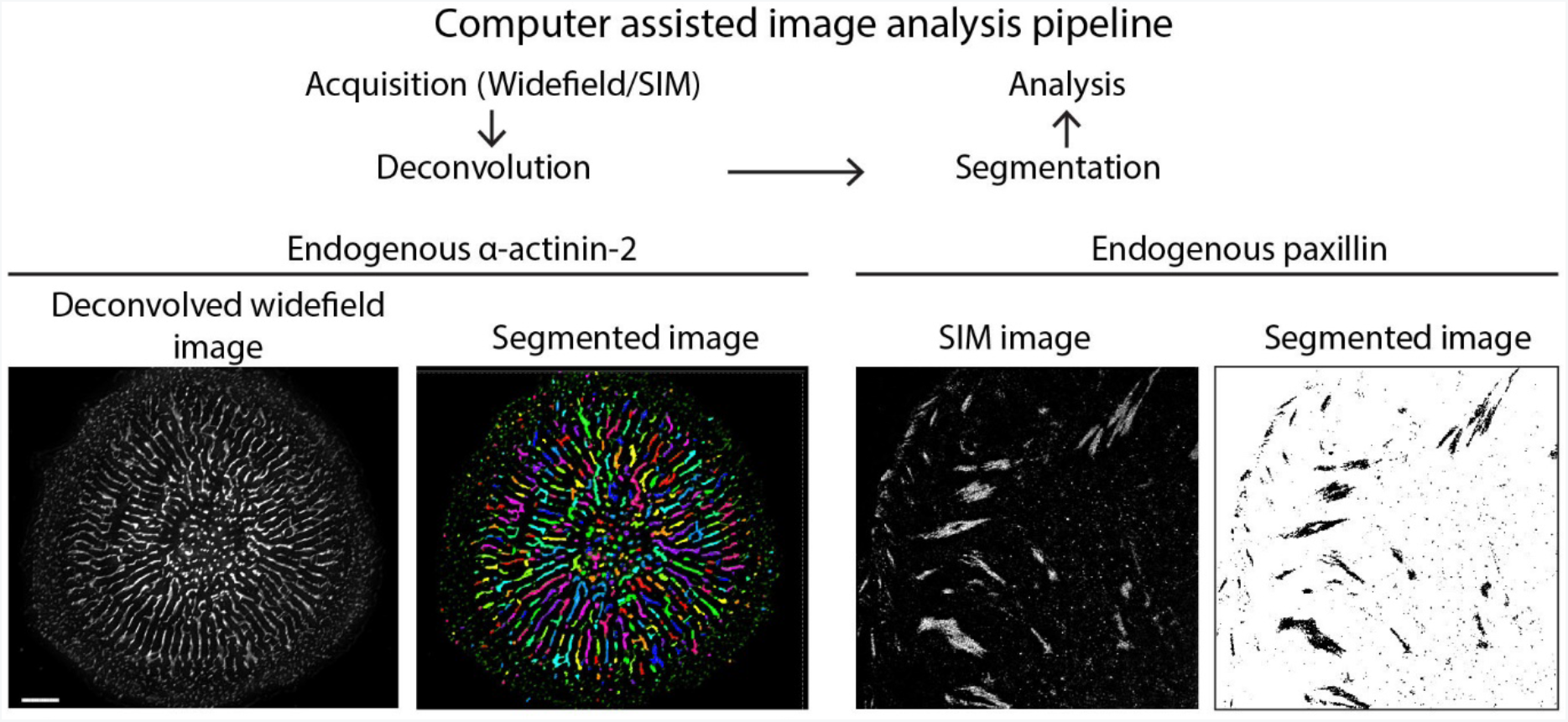
Computer assisted image analysis pipeline. For measurement of a-actinin-2 Z-line lengths, widefield fluorescence image stacks (gray) were deconvolved and segmented using Nikon Elements Software (colorized binary file). The long axis lengths of Z-bodies were exported to MATLAB and data was filtered and processed to generate median Z-line measurements. For paxillin quantification, SIM micrographs were used for thresholding in FIJI. Filtering and further analysis was performed in MATLAB. Scale bar: 5 µm.

**Supplementary Figure 3.**
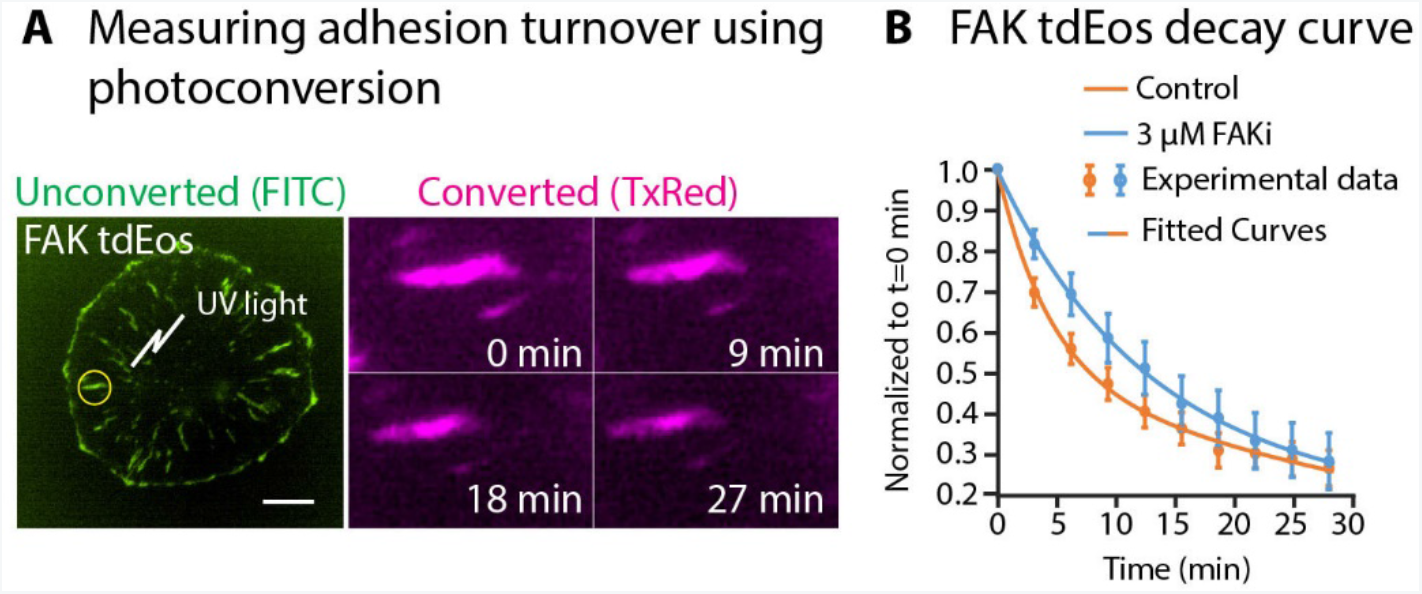
Measurement of adhesion dynamics using photoconversion. A) hCM expressing FAK tdEos. Yellow ROI was used to photoconvert a focal adhesion (green) and then followed in the converted channel (magenta) for 27 min. B) Averaged decay curve for FAK tdEos in untreated control versus 3 µM PF-228 hCMs. Curve fitting was performed in MATLAB to generate half-life and mobile fractions. Scale bar: 10 µm.

